# In Vivo AGO-APP For Cell-Type- and Compartment-Specific miRNA Profiling in the Mouse Brain

**DOI:** 10.1101/2025.08.07.669091

**Authors:** Surbhi Kapoor, Andrea Erni, Gunter Meister, Harold Cremer, Christophe Beclin

**Affiliations:** Aix-Marseille Université, Centre National pour la Recherche Scientifique (CNRS), Institut de Biologie du Développement de Marseille (IBDM), Marseille, France; Regensburg, Center for Biochemistry (RCB), University of Regensburg, Regensburg, Germany1

**Keywords:** MiRNA, Argonaute, cortical glutamatergic neurons, olfactory bulb interneurons, Dicer, TNRC6, T6B, post-synapse

## Abstract

AGO-APP through the expression of the T6B peptide permits the isolation of Ago-bound miRNAs. Here we present the generation and characterization of two transgenic mouse lines allowing to perform AGO-APP in vivo. First, we generated mice for CRE-dependent expression of T6B in the cytoplasm. Using this line we performed Ago-APP in olfactory bulb (OB) inhibitory interneurons and cerebral cortex excitatory neurons. Bioinformatic analysis validated the high reproducibility of the approach. It also demonstrated that, despite global miRNome conservation between the two cell types, a set of miRNAs including the miR-200 family and the miR-183/96/182 cluster, is massively enriched in OB interneurons which is consistent with previous observations. In the second mouse line T6B is fused to the postsynaptic protein PSD95. Isolation of T6B-PSD95 fractions from OB and cortical neurons identified specific sets of post-synapse enriched miRNAs. Gene ontology analyses confirmed that these miRNAs preferentially target mRNAs related to synaptic functions.

**Motivation:** Collectively, miRNAs are required for all biological processes in eukaryotes. However, due to a lack of appropriate methodological tools, the actual mode of action of individual miRNAs in a given cell is currently poorly understood. In this article, we describe a novel approach to study the specific expression of miRNAs in vivo, at the level of cell type and in a specific cell compartment.

## Introduction

MicroRNAs (miRNAs) are small non-coding RNA molecules that regulate gene expression by binding to complementary sequences on target mRNAs. The exact number of miRNAs expressed by the mammalian genomes is still debated. However, recent estimates set this number at around 2300 in humans^1^ and 2200 in mice^2^. Interference with the function of the miRNA pathway, for example through genetic inactivation of the miRNA-generating ribonuclease Dicer, induces severe defects in animal development and tissue homeostasis, demonstrating the important physiological role of this process^3,4^.

MiRNA/mRNA interactions are highly promiscuous. Indeed, a single miRNA can bind to many mRNAs and a specific mRNA can present binding sites for several miRNAs leading to the prediction that more than 60% of transcripts are miRNA targets^5^. Therefore, detailed information about the expression of miRNAs and mRNAs in a given cell type is essential to infer their functional interactions. For mRNAs such expression information is nowadays easily accessible, at the tissue and even at the single cell levels, through the development of single-cell RNA sequencing protocols or spatial transcriptomics. In contrast, the analysis of miRNA expression is still highly challenging, especially when complex tissues or rare cell types need to be studied in vivo.

Current protocols for miRNA expression analyses depend on relatively large amounts of input material, and single cell sequencing is still highly experimental^6,7^. Imaging approaches to miRNAs, for example by in situ hybridization, are problematic due to their small size and the high degree of promiscuity^8^. FACS sorting of dissociated primary cells before sequencing has been used to address cell type specific expression, but depends on the availability of suitable markers^9^. Moreover, morphologically complex cell types, like for example mature neurons, are highly refractory to intact dissociation. Finally, there is evidence that not all expressed miRNAs in a given cell are bound to Argonaute (AGO) protein and integrated into an RNA-induced silencing complex (RISC), indicating that only a fraction of all expressed miRNAs is active in inhibition^10–12^.

In addition to such considerations concerning miRNAs that are active in the cytoplasm, there is evidence that miRNAs have specialized roles when targeted to specific cellular compartments. For example, miRNAs were proposed to act in axons^13,14^ or the nucleus^15–17^ . However, the specific presence and function of miRNAs is best documented for the post-synapse, for which several studies suggest a role in regulating local translation, and consequently synaptic plasticity^18–20^.

In light of this high degree of expression and functional complexity it is evident that new technologies and approaches are needed to overcome the existing roadblocks. T6B is a small peptide of 80 amino acids (aa) derived from the human TNRC6B protein which itself is part of the RISC, where it links the AGO-miRNA-mRNA complex with the downstream mRNA degradation machinery^21^. Previous work showed that although T6B is non-functional, it retains the ability to bind all different AGO proteins involved in the miRNA pathway, and can be used in vitro to capture AGO-bound miRNAs in pull-down experiments^22^ in a process called AGO-APP. Based on this initial finding we showed recently that transgenic expression of T6B in Drosophila melanogaster allows the isolation and analysis of miRNAs from neural stem cells, neurons and glial cells with high precision. This approach led to the discovery of a regulatory module of miRNAs that cooperatively preserved neural progenitors from premature differentiation^23^ .

We present two transgenic mouse lines allowing the CRE-dependent expression of T6B, either in the cytoplasm or specifically targeted to the post-synapse. We use these lines for the direct and reliable isolation and analysis of miRNAs from specific neuron types in the vertebrate brain and provide proof-of principle that the subcellular localization of miRNAs at the post-synapse can be addressed by this approach.

## Results

### Sustained expression of T6B in neuronal progenitors does not perturb postnatal neurogenesis

During postnatal neurogenesis in the rodent forebrain neural progenitors (NPs) located in the subventricular zone (SVZ) generate permanently neuronal precursors that migrate into the olfactory bulb (OB). Here they mature as GABAergic interneurons, granule cells and periglomerular neurons, and form functional synapses around five weeks after their birth^24^. This system can be easily manipulated by postnatal in vivo electroporation of the ventricular wall^25,26^, and phenotypic consequences can be studied at high resolution, making it a model of choice for the investigation of molecular events underlying neuron generation. Moreover, ongoing neurogenesis in the SVZ-OB neurogenic system was shown to implicate the function of specific miRNAs, acting at different steps of the process. For example, miR-124 favors the exit from a neural progenitor state^27^ whereas miR-9 controls subsequent proliferation^28^. MiR-7^29^ and miR-124^30^ regulate fate determination, while miR-200^31^ and miR-132^32^ participate in final maturation of new neurons.

We investigated the level of dependency of this neurogenic system on a functional miRNA pathway using a conditional dicer mutant mouse line (Dicer-floxed;^33^). Dicer is a key enzyme in the generation of miRNAs and its lack leads to complete inactivation of this regulatory system ^4,34^. Postnatal in vivo electroporation was used to express CRE in NPs along the ventricular walls of mice carrying a conditional mutation in the Dicer gene. To visualize recombined cells by RFP expression, we introduced the CRE-dependent reporter allele Ai14 (B6.Cg-*Gt(ROSA)26Sor^tm^*^14^*^(CAG-tdTomato)Hze^*/J ; JAX stock #007914 ;^35^). Electroporation was targeted to the lateral ventricular wall, that generates predominantly OB granule cells.

During the first two weeks after electroporation comparable numbers of RFP positive neurons were detected in the OB granule cell layer of control (CRE:+/fl) and dicer-deficient (CRE:fl/fl) animals (Fig. 1A). Over the following weeks the number of RFP positive neurons almost doubled in control mice due to ongoing neuron production by recombined NPs in the SVZ. In contrast, there was a significant drop in the density of labelled neurons in dicer -/- animals, until the almost total loss of all RFP positive cells at 7 weeks post electroporation (Fig. 1A). We conclude that a functional miRNA pathway is essential for the generation of new neurons in the SVZ-OB system.

**Figure 1.**
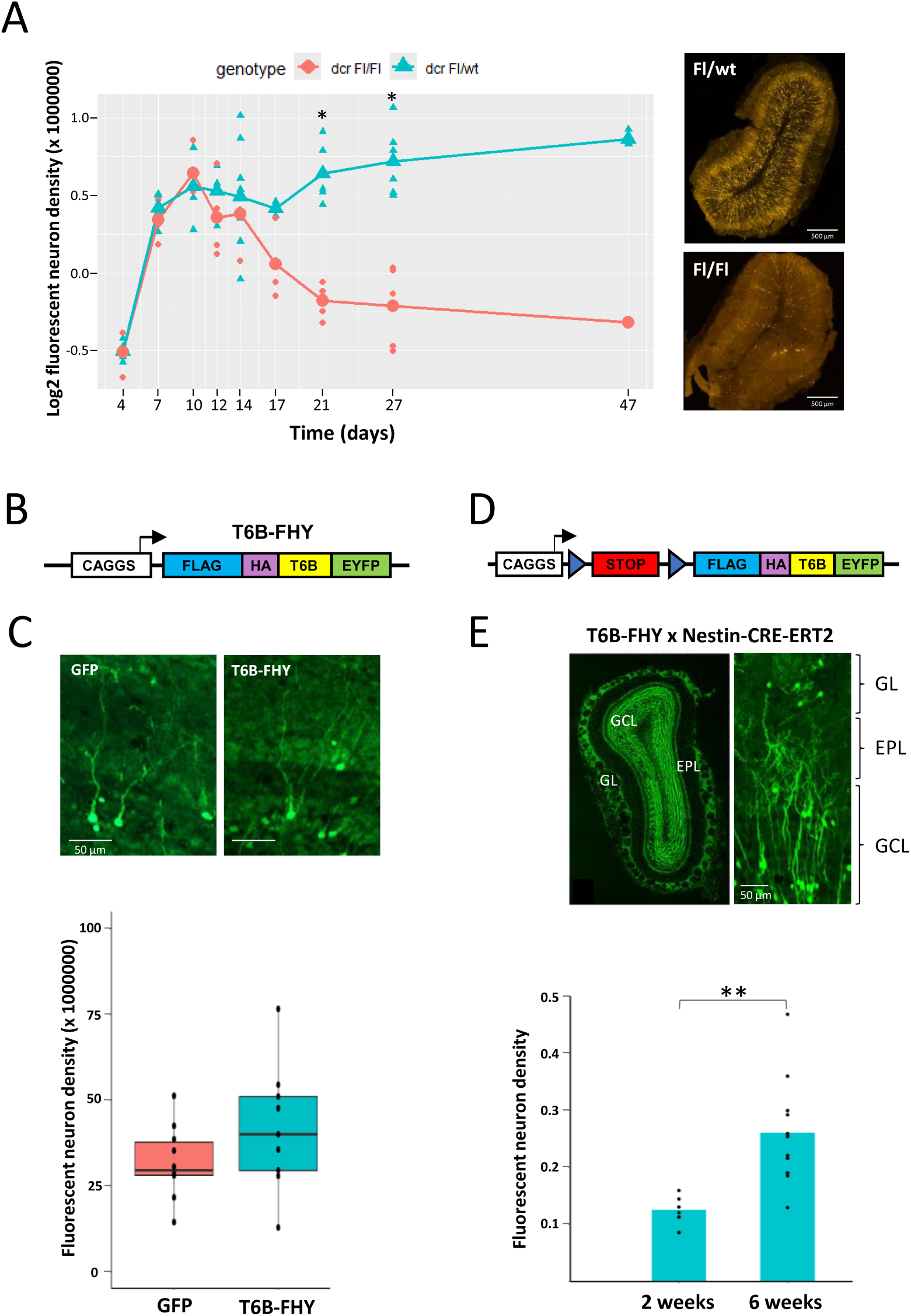
Sustained expression of T6B in neuronal progenitors does not perturb postnatal neurogenesis. (A). Left: Time course of the density (log2 scale) of recombined neurons in OB sections after CRE electroporation in the SVZ of P1 animals heterozygous (green) or homozygous (red) for a conditional mutant allele of dicer. Cell density for each individual is shown as red circles or green triangles. For each individual 3 to 4 bulb sections were quantified. *p-value ≤ 0.05. Right: Representative mosaic images (stitched) of OB coronal section from dicer heterozygous (Fl/wt) or homozygous (Fl/Fl) animals. (B) Scheme of the pCX-T6B-FHY expression plasmid. (C) Representative images stained with anti-GFP antibody showing recombined OB neurons 5 weeks after electroporation at P1 using pCX-GFP (left) or pCX-T6B-FHY (right) plasmid. Box plot comparison of the density of fluorescent neurons per OB section (multiplied by 10^6^) after electroporation using the pCX-GFP control plasmid (red) or the pCX-T6B-FHY plasmid (green). Cell density in individual animals is shown as a black dot. For each individual 3 to 4 bulb sections were quantified. (D) Schematic representation of the construct used to generate the conditional T6B-FHY expressing transgenic mouse line. LoxP sites, encompassing a STOP cassette, are indicated as triangles. (E) Top: Representative mosaic image (stitched) of a whole OB section stained with anti-GFP antibody 5 weeks after tamoxifen injection in a P1 T6B-FHY transgenic animal (left). High magnification (right) showing individual T6B-FHY expressing neurons in the granule cell layer (GCL) and the glomerular layer (GL). EPL: external plexiform layer. Bottom: Density of fluorescent neurons in OB sections of T6B-FHY animals at 2 and 5 weeks after tamoxifen injection. Six (2 weeks) and 12 (5 weeks) animals were treated. Three OB sections were quantified per animal. **p-value ≤ 0.01.

Previous in vitro work provided evidence that binding of T6B to AGO can disrupt the RISC complex and interfere with the function of the miRNA pathway^36^. In vivo, de-repression of several target mRNAs and defects in homeostasis of skeletal muscles and heart have been observed. However, changes in miRNA levels were not detected in these studies^37,38^. In light of these findings, we investigated if T6B expression impacts on OB neurogenesis.

A pCX-T6B-FHY expression plasmid derived from the pCX-mcs2 vector^39^ and containing the T6B coding sequence fused to FLAG and HA tags, as well as the fluorescent protein EYFP (henceforth called T6B-FHY), under the control of the ubiquitous CAGGS promoter (Fig. 1B), was introduced into NPs by postnatal electroporation at P1. Analyses of EYFP positive neurons in the OB at 5 weeks post-electroporation did not reveal obvious morphological alterations or significant differences in the number of granule cells in comparison to controls in which NP were electroporated with a pCX-GFP plasmid^39^ (Fig. 1C), indicating that any potential T6B-induced perturbation was too minor to affect the proper generation and integration of new cells.

Based on these observations we generated a transgenic mouse line allowing the CRE-dependent expression of T6B-FHY (Fig. 1D) under the CAGGS promoter. The construct was integrated into the Rosa26 locus^40^ by homologous recombination. Resulting transgenic mice were bred to a Nestin-CRE-ERT2 transgenic line^41^ to drive temporal controlled recombination in postnatal SVZ NPs upon tamoxifen injection at P1. Two weeks after induction, the granule cell layer and the glomerular layer (GL) of the OB were strongly colonized by EYFP positive cells with the typical morphology of young granule cells and periglomerular neurons (Fig. 1E). At five weeks post electroporation the density of newborn neurons more than doubled (Fig. 1E), comparable to the increase observed in Dicer +/fl mice and strikingly different from the cell loss observed in Dicer mutants (Fig. 1A). Thus, other than after total loss of the miRNA pathway induced by Dicer KO (Fig. 1A), neurogenesis in the SVZ-OB system proceeds at normal levels in the presence of T6B-FHY expression, indicating that any potential interference with miRNA activity is minor, not obviously impacting neuronal integration and survival.

#### miRNA isolation from brain tissue by vivo AGO-APP

Binding of T6B to AGO disrupts RISC by displacing TNRC proteins from the complex (Fig. 2A). Our previous work in drosophila melanogaster demonstrated that the miRNA:AGO:T6B-FHY complex can be efficiently pulled down with anti-GFP nanobodies^23^. Subsequently, miRNA is released and can be analyzed by sequencing. This approach was used to generate a high-resolution map of miRNA expression in drosophila neural stem cells, neurons and glial cells^23^.

**Figure 2.**
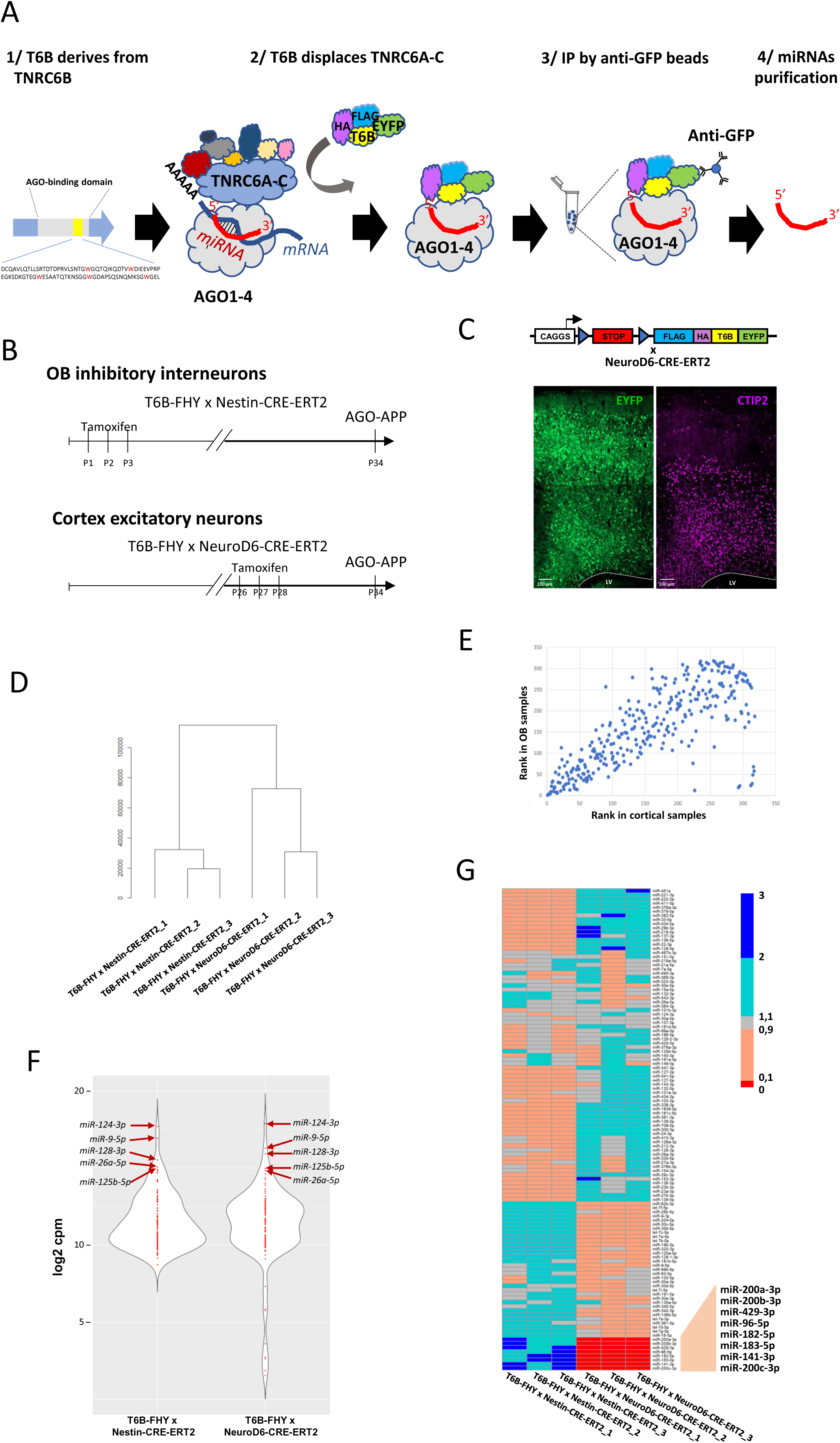
miRNA isolation from brain tissue by vivo AGO-APP. (A). Schematic representation of the T6B peptide and the AGO-APP procedure. (B) Time line of the experiments aimed at generating AGO-APP samples from both OB inhibitory interneurons and cortical excitatory projection neurons. (C) Mosaic image (stitched) showing that tamoxifen injection into adult animals derived from the crossing of the T6B-FHY with the NeuroD6-CRE-ERT2 transgenic mouse lines results in EYFP expression in cortical glutamatergic excitatory neurons across all layers of the motor cortex (left). CTIP2 staining (red) validates expression in deep layer cortical neurons (layers V and VI). s. (D) Hierarchical clustering of miRNA sequencing of AGO-APP samples purified from T6B-FHY x Nestin-CRE-ERT2 and T6B-FHY x NeuroD6-CRE-ERT2 animals shows that neurons of a given type tightly cluster. (E) Spearman correlation between the mean of the cpm reads of the three T6B-FHY x Nestin-CRE-ERT2 samples (Y-axis) and the three T6B-FHY x NeuroD6-CRE-ERT2 samples (X-axis). Each dot represents one miRNA among all miRNAs that were expressed at a significant level (sum of cpm reads of all 6 samples greater than 100), r = 0,7565295, adjusted p-value < 2.2 e-16. (F) Violin plots showing in OB interneurons or cortical neurons the log2 of the mean per condition of miRNAs normalised expression (cpm). Only miRNAs whose sum of cpm reads from the 6 samples is greater than 5000 are shown. The 5 most highly expressed miRNAs in each condition are labelled. (G) Heat map of normalized values (computed by dividing the normalized cpm read counts by the mean of all samples) for all miRNAs with a sum of cpm reads of all 6 samples greater than 5000. A close-up is shown for the 8 miRNAs showing a striking overexpression in OB (T6B-FHY x Nestin-CRE-ERT2) versus cortical neuron (T6B-FHY x NeuroD6-CRE-ERT2) samples.

We aimed at using this new approach to provide insight into the miRNA expression in different types of vertebrate neurons. Indeed, due to their complex morphology and their tight contact with other cells, isolation of intact neurons from vertebrate brain tissue, and analysis of their cytoplasmic content, is still highly challenging^42^. As a consequence, information about neuronal subtype specific expression of miRNAs is currently limited. AGO-APP is suited to overcome this problem as T6B-FHY can be expressed under the control of suited promoters in specific cell types for directly isolating miRNAs from tissue homogenates, without the need for cell isolation. We used this approach to compare miRNA expression between GABAergic inhibitory neurons and glutamatergic excitatory neurons.

To isolate miRNAs from GABAergic neurons we used the above-described approach based on induction of CRE recombination in SVZ progenitors by the Nestin-CRE-ERT2 driver, subsequently leading to T6B-FHY expression in inhibitory OB interneurons (Fig. 1D,E). Three tamoxifen injections were performed at P1, P2 and P3 to induce T6B-FHY in a large number of NPs (Fig. 2B). Five weeks later, when newborn neurons derived from these NPs arrived in the OB and were fully mature, OBs were dissected, homogenized and miRNAs were isolated by AGO-APP.

To drive expression of T6B-FHY in mature glutamatergic projection neurons we used a well characterized inducible NeuroD6-CRE-ERT2 line, that drives CRE recombination into cortical excitatory neurons, including layer V and VI neurons expressing CTIP2 (Fig. 2C^43^). For AGO-APP, tamoxifen was injected three times at P28-P30 and cortical tissue containing mature T6B-FHY expressing neurons was harvested four days later (Fig.2B).

Subsequently, miRNAs pulled down by the AGO-APP procedure were sequenced using a modified version of the AQ-seq protocol^44^, optimized for samples with low RNA concentration. Hierarchical clustering of samples based on miRNA sequencing reads showed that the 3 OB replicates and the 3 samples issued from cortex clustered tightly together respectively (Fig. 2D), demonstrating high reproducibility of the approach.

After normalization to ’counts per million’ (cpm) and the exclusion of miRNAs poorly expressed over all samples (sum of cpm reads from all samples less than 100) we compared the global miRNA expression pattern between the two cell types using Spearman rank correlation (Fig. 2E). This analysis of correlation between miRNA expression in both neuron types resulted in a r coefficient of 0,7565295 and an associated p-value lower than 2.2 e-16, indicating an overall conservation of the miRNomes. This overlap is well exemplified by the observation that the 5 miRNAs showing strongest expression in GABAergic neurons were also the highest present in glutamatergic neurons (miR-124-3p, miR-9-5p, miR-128-3p, miR-125b-5p, miR-26a-5p) (Fig. 2F).

However, despite obvious overlap, the cluster analysis suggested differences in miRNA content from both neuron types (Fig. 2D). For a graphical comparison of both miRNomes we built a heat map focusing on the 117 most strongly expressed miRNAs (sum of cpm reads from all samples greater than 5000). Heat map representation of the homogenized data allowed to pinpoint a set of eight miRNAs, belonging either to the miR-200 family (miR-200a-3p, miR-200b-3p, miR-429-3p, miR-141-5p, miR-200c-3p) or to the miR-183/96/182 cluster (Fig. 2G), that showed striking differential expression, with strong presence in OB neurons but quasi absence from cortical neurons. This observation was fully consistent with our previous findings in raw OB tissue^31^, further validating the AGO-APP approach. Moreover, these eight miRNAs have been functionally implicated in the control of neurogenic events. Indeed, the miR-200 family has been shown to be involved in the maturation of OB interneurons and the development of olfactory sensory neurons^45,46^, while the miR-183/96/182 cluster controls maturation and maintenance of retinal photoreceptors^47,48^.

We then performed a DESeq2 statistical analysis to systematically determine differentially expressed miRNAs between the two types of neurons. As expected, strongest differential expression was observed for the above-mentioned set of eight miRNAs over-expressed in the OB (Table 1). Moreover, we identified a total of 209 miRNAs for which the padj value was lower than 1%. Implementing a fold change threshold of 2 and a base mean of 500, this number was reduced to 62, among which 30 were found over-expressed in cortical neurons (Table 1). Importantly, this list of cortex-enriched miRNAs comprised miR873a-5p, miR218-5p, miR137-3p, miR27-3p, miR-185-5p and miR-433-3p, six miRNAs for which previous GWAS analyses showed an association with neurological disorders involving the cortex, such as autism spectrum disorder^49^, schizophrenia^50–52^, bipolar disorder^53,54^ or depressive symptoms^55^.

Overall, these data show that in vivo AGO-APP using the new transgenic mouse line is highly reproducible and suited for determining miRNA expression in specific neuron subtypes at high resolution.

#### Analyzing miRNA expression at the post-synapse

The above-described experiments demonstrate that the expression of the T6B peptide in transgenic mice allows the capture of Argonaute-bound miRNAs from the entire cytoplasm of different neuron types. However, besides their well described role in general mRNA translational control, there is strong evidence that miRNAs have specialized roles when targeted to specific cell compartments, like for example the post-synaptic aspect of glutamatergic synapses^18–20^.

In mammals, post-synaptic density protein 95 (PSD95) is confined to the postsynaptic compartment of excitatory synapses^56^ and has been used to efficiently transport fused protein to the post-synapse, without altering synaptic morphology or physiology^57^. Based on these properties we generated a second transgenic line, in which T6B-FHY was fused to PSD95 (Fig. 3A). First, we studied subcellular localization, in either OB inhibitory neurons with the Nestin-CRE-ERT2 driver, or in cortical excitatory neurons, using NeuroD6-CRE-ERT2 (Fig. 3B). Analyses of EYFP fluorescence at p34 in both lines validated that PSD95-T6B-FHY showed always the striking punctate staining typical of synaptic localization^57^, whereas labelling in cell bodies (surrounded by a dotted line) was very faint, different from the diffuse but strong cytoplasmic labelling observed for T6B-FHY alone (Fig. 3C, but see Fig. 1E). PSD95 driven localization at post-synapse was validated using electron microscopy of immunogold stained material from OB and cortex, that showed clusters of electron dense particles mainly confined to post-synaptic structures (Fig. 3D)

**Figure 3.**
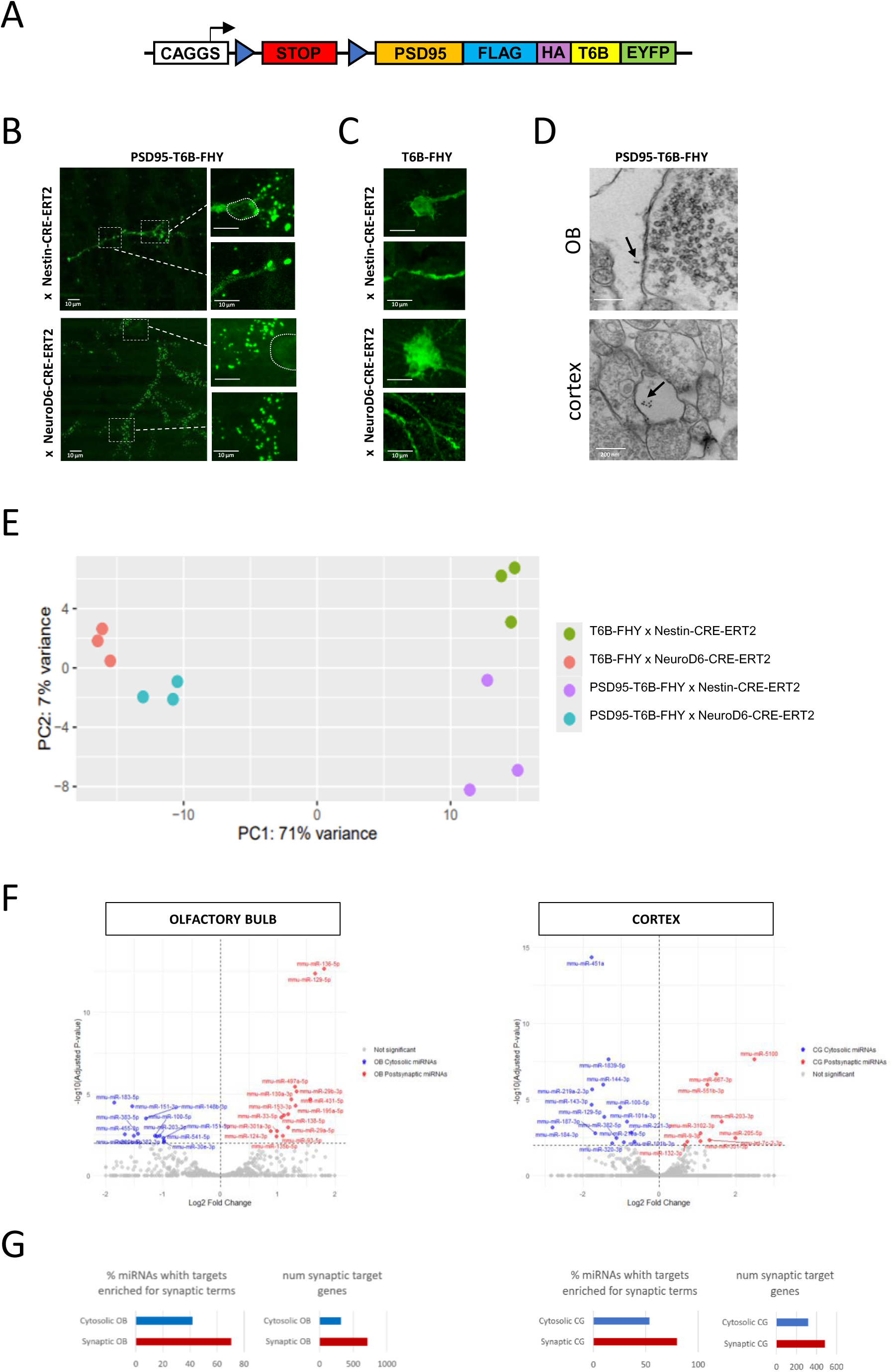
In vivo AGO-APP to analyze miRNA expression at the post-synapse. (A) Schematic representation of the transgene used to generate the PSD95-T6B-FHY expressing mouse line. (B) Confocal images showing olfactory bulb interneurons (top) and cortical glutamatergic projection neurons (bottom) expressing PSD95-T6B-FHY. Sections were labelled using an anti-GFP antibody. Close-ups point to representative cell body (top) and dendrite segment (bottom) showing punctate staining in the periphery and quasi absence in the cytoplasm of both cell compartments. (C) Representative images of OB interneurons (top) and cortical glutamatergic projection neurons (bottom) expressing T6B-FHY without PSD95. in both cell types, the staining is homogeneous, lacking the punctate aspect indicative of synaptic localization, in the cell body (top) or within dendrite (bottom). (D) Electron microscopy images after anti-GFP gold bead labelling of synapses of OB interneurons (top) or cortical projection neurons (bottom) expressing PSD95-T6B-FHY. Arrows show groups of beads in the postsynaptic compartment. (E) PCA analysis of miRNA sequencing data after AGO-APP performed on the 4 transgenic lines used in the study (3 replicates per condition): T6B-FHY x Nestin-CRE-ERT2, T6B-FHY x NeuroD6-CRE-ERT2, PSD95-T6B-FHY x Nestin-CRE-ERT2, PSD95-T6B-FHY x NeuroD6-CRE-ERT2. (F) Volcano plot for DESeq2 results showing miRNAs differentially present at the post-synapse in OB interneurons (left) and cortical projection neurons (right). MiRNA enriched at the post-synapse or in the cytosol are colored in red and blue, respectively. (G) GO analysis of predicted mRNA targets of the miRNAs that were found differentially targeted to the post-synapse in olfactory bulb interneurons (left) and cortical glutamatergic projection neurons (right). Two parameters were quantified for both types of neurons. Left panel: percentage of miRNAs that are enriched (red) from the post-synapse and for which the list of Targetscan predicted mRNA targets are significantly enriched in synaptic genes. Right panel: Cumulative amount of mRNA targets that are predicted to be synaptic.

Next, we performed AGO-APP and small RNA sequencing on either OB or cortical tissue dissected from the transgenic mouse lines expressing PSD95-T6B-FHY and compared the resulting data with those previously obtained with the T6B-FHY line. PCA analyses demonstrated strong clustering of all biological replicates (3 per condition) underlining the high reproducibility of the approach (Fig. 3E).

Neuron type explained most of the observed variability, whereas variability explained by cell compartment (cytosol versus post-synapse) was considerably lower. For both neuron types, analyses of differential expression between T6B-FHY and PSD95-T6B-FHY isolates identified small sets of miRNAs that showed significantly different expression levels between both compartments (Fig. 3F) indicating preferential post-synaptic enrichment of a subset of miRNAs. Interestingly, 6 among 15 miRNAs enriched in the OB post-synaptic fraction (miR-138^58^, miR-153^59^, miR-135^60^, miR-124^61^, miR-29a/b^62^ and miR-129-5p^63^ and 2 among the 10 cortical post-synaptic high miRNAs (miR-9^58^ and miR-132^64^) have been previously associated to synaptic function in other studies. Finally, we predicted potential target mRNAs of the miRNAs enriched in the OB and cortical PSD95-T6B-FHY fractions, using the Targetscan algorithm. Gene ontology analyses of this dataset revealed that the AGO-APP postsynaptic fractions of OB and cortex neurons were enriched for miRNAs targeting mRNAs that are linked to synaptic terms (Fig. 3G).

Altogether, this data provides strong evidence that PSD95-T6B-FHY is a potent tool for the isolation of postsynaptic miRNAs.

#### Limitations of the study

Whereas our control experiments do not reveal perturbation of neurogenesis by T6B expression, it cannot be excluded that T6B expression reduces slightly the level of miRNA regulation and consequently modifies marginally the expression of a subset of genes.

In this work, we applied AGO-APP to decipher the differential expression of Ago-bound miRNA between OB GABAergic and cortical glutamatergic neurons, and also between the cytosol and the post-synaptic compartments of these 2 neuron types. Although these data are consistent with previous findings, we do not provide an in-depth functional analysis. Indeed, such direct experimental validations are difficult to design, especially for validating the enrichment of miRNAs at the post-synapse of specific neurons under in vivo conditions. Alternatively, indirect validation may be envisaged by, for example, correlating the post-synaptic enrichment of miRNAs presumably involved in plasticity with artificial change in neuronal activity.

The AGO-APP technology presented in this manuscript allows the immunoprecipitation of AGO bound miRNAs, as does the AGO-CLIP technology. As demonstrated in this manuscript AGO-APP presents several advantages over the concurrent AGO-CLIP to analyze in detail the activity of Ago-bound miRNAs in the cell, like for example the possibility to analyze the targeting of miRNAs to specific compartments of the cell. However, although we did not specifically address this point, it is likely that the displacement of TNRC6 protein by T6B will weaken the miRNA-mRNA association in the cell^65^.

Therefore, it might be more challenging to co-isolate miRNAs together with their target transcripts.

## Discussion

MiRNAs emerged as important regulators of gene expression in the cell. Collectively they are required for most, if not all, cellular processes, as demonstrated by the severe phenotypic alterations that are induced when the entire regulatory pathway is inactivated^3^. Our finding that Dicer deficiency is not compatible with the generation of new interneurons in the postnatal OB represents a further display of this essential role.

However, beyond the general conclusion that the miRNA pathway is required for a process to occur, the precise mechanism by which the miRNAs regulate is still largely unclear, mainly due to conceptual and technical limitations. Indeed, experiments addressing the role of specific miRNAs generally induce only minor phenotypes^66–68^, not compatible with an all-or-nothing function of individual miRNA-target interactions. In agreement, several recent studies provided evidence that entire sets of miRNAs cooperate to control the expression of a particular gene or a regulatory network^23,68^.

Technically, analyzing the expression and function of miRNA at the cellular level is highly challenging. For example, the amount of needed input material is currently not compatible with routine single cell sequencing, and histological approaches are still challenging. Moreover, cell isolation in vivo depends generally on tissue microdissection and dissociation followed by cell sorting. Disruption of complex cells like neurons during dissociation will lead to miRNA loss. Finally, functional studies by inhibiting the expression of specific miRNAs are often complicated by the fact that many miRNA loci are located within protein coding genes or produced from more than one genomic location^69^.

The situation is even more complicated. Indeed, it has been reported that not all miRNA expressed in a given cell are involved in regulation, but that “silent” and “active” miRNAs coexist at a given time point^10–12^. Thus, to launch subsequently complex functional studies, simply isolating and analyzing the entire complement of cellular miRNAs might not provide sufficiently conclusive information and may lead to the prediction of multiple non-functional interactions. Finally, there is considerable data indicating miRNAs transport to, and function in, specific subcellular compartments like axons, nucleus or synapse. Unbiased investigation of miRNAs showing such subcellular localization in vivo is currently impossible.

In vivo AGO-APP represents a tool to overcome many of these limitations. First, transgenic expression of T6B allows the isolation of miRNAs from complex cellular environments like brain tissue. Due to the expression of T6B as a fluorescent fusion protein, cell types to be analyzed can be studied at high resolution, allowing the correlation of morphological and sequencing data. Also, as the miRNA-AGO-T6B complex forms in the intact cell, disruption of the cytoplasm does not lead to loss of material but is an intrinsic part of the isolation approach. Finally, in vivo-AGO-APP is a benchtop approach, not requiring sophisticated multistep manipulations, specialized equipment or expertise in FACS, allowing the recovery of sufficient amounts of miRNAs for reliable sequence analysis leading to meaningful data.

AGO-APP is based on immunoprecipitation of AGO-miRNA complexes and therefore related to the well-established AGO-CLIP technology. Like AGO-CLIP, AGO-APP isolates only AGO bound miRNAs that are considered to be the “active fraction” in a background of all expressed miRNAs^11^. However, other than AGO-CLIP, AGO-APP is based on the expression of an exogeneous peptide that can be controlled by transgenic technology in vivo. This allows the determination of cell type specific miRNomes with unpreceded precision. Here we isolated miRNAs from two spatially separated cell populations, cortical excitatory neurons and OB interneurons. We find overall similarity between the ago-bound miRNA pools, suggesting global conservation of the regulatory pathway in neurons.

However, the high resolution of AGO-APP permitted to resolve quantitative variations in miRNA numbers between both cell types and also led to the identification of an OB specific subset of miRNAs that might act in concert to fulfill specific roles in these neurons.

Finally, our approach allows to systematically address the presence of miRNAs in a cellular sub-compartment. Indeed, there is a vast literature describing post-synaptic miRNAs and their specific activities during homeostasis and plasticity^20^. Addressing such functions in a physiologic context is challenging, if not impossible, with the available technologies. However, as we show here, T6B can be specially targeted to the post-synapse and allows the isolation and analyses of miRNAs enriched in this compartment. We found that miRNAs enriched at the post-synapse, as identified by quantifying the ratio between cytoplasmic and postsynaptic miRNAs, are very different between the two cell types analyzed, despite an overall conservation of both miRNomes. Moreover, the list of post-synaptic miRNAs we identified in this study exhibits only very partial overlap with the synaptic miRNAs reported in the literature. Thus, the list of miRNAs preferentially targeted to the post-synapse seems to be cell-type specific suggesting that the post-synapse targeting of miRNA is not an intrinsic property of miRNAs but instead is dependent on the physiological environment.

AGO-APP represents a technical advance in understanding the expression and mode of action of miRNA. Using two different CRE drivers, this technology allowed us to compare the expression of AGO-bound, and therefore actively inhibiting, miRNAs in two given neuron types. By selecting the appropriate CRE driver, the scope of this technology can be extended to any cell type of the brain and other tissues. We have also shown that AGO-APP can be used to study the enrichment of specific miRNAs at the post-synapse in living animals. Therefore, AGO-APP will allow to study the local binding of synaptic miRNAs to AGO as a function of neuronal activity and plasticity, for example during learning and memory. Finally, by using alternative molecular tags, other than PSD95, it is conceivable that AGO-APP could be used to study miRNA activity in other cell compartments for which local functions have been suggested, such as the axon or the nucleus.

## Resource availability Lead contact

Requests for further information, resources, and reagents should be directed to and will be fulfilled by the lead contact, Christophe Beclin.

## Materials availability

Plasmids and strains generated in this study are available upon request from the lead contact, Christophe Beclin.

## Data and code availability

The sequencing datasets generated and analyzed during this study have been submitted to the functional genomics data collection ArrayExpress and have received the accession number E-MTAB-15124.

Scripts used to generate the figures presented in this paper are available from the lead contact upon request. This study does not report original code.

Any additional information required to reproduce this work is available from the lead contact.

## Acknowledgments

The Cremer lab has been supported by the ANR (MicroRNAct, ANR-17-CE16-0025; Uncoding, ANR-21-CE16-0034; Miniature, ANR-21-CE13-0003) and the Fondation pour la Recherche Medicale (Program Equipe FRM). Surbhi received a PhD fellowship from Neuroschool Marseille from Aix Marseille University. Andrea Erni received a postdoc fellowship from the Swiss National Science Foundation (SNSF).

We thank Lena Vilvandre for genotyping of the animals. We thank Marie-Catherine Tiveron for helping in confocal image acquisition and Nathalie Core for critical reading of the manuscript. We also thank the imaging facility at IBDM, member of the National Infrastructure France-BioImaging (https://ror.org/01y7vt929) supported by the French National Research Agency (ANR-24-INSB-0005 FBI BIOGEN) as well as the animal facilities. The electron microscopy experiments were performed on the PiCSL-FBI core facilty (IBDM, AMU-Marseille), member of the France-BioImaging national research infrastructure (ANR-10-INBS-04) and member of the Marseille Imaging Institute, an Excellence Initiative of Aix Marseille University A*MIDEX, a French “Investissements d’Avenir” programme (AMX 19 IET 002). The funders had no role in study design, data collection and analysis, decision to publish, or preparation of the manuscript. We thank members of the Meister lab for providing the T6B-FHY construct and for helpful advice.

## Author contributions

C.B. and H.C. designed the study and wrote the paper. SK and AE performed the experiments. GM helped in designing the in vivo Ago-APP protocol.

## STAR★Methods

### Key resources table

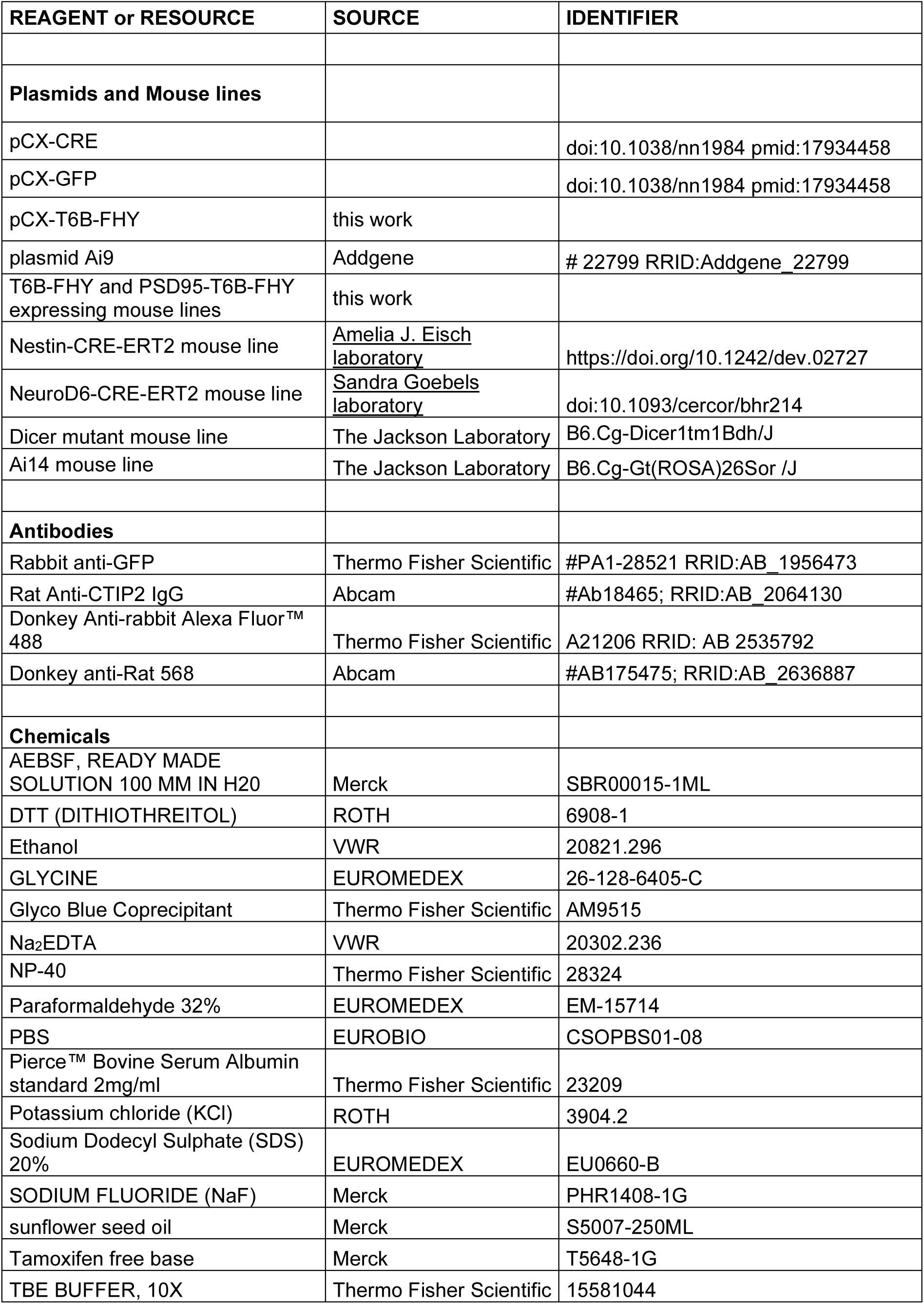
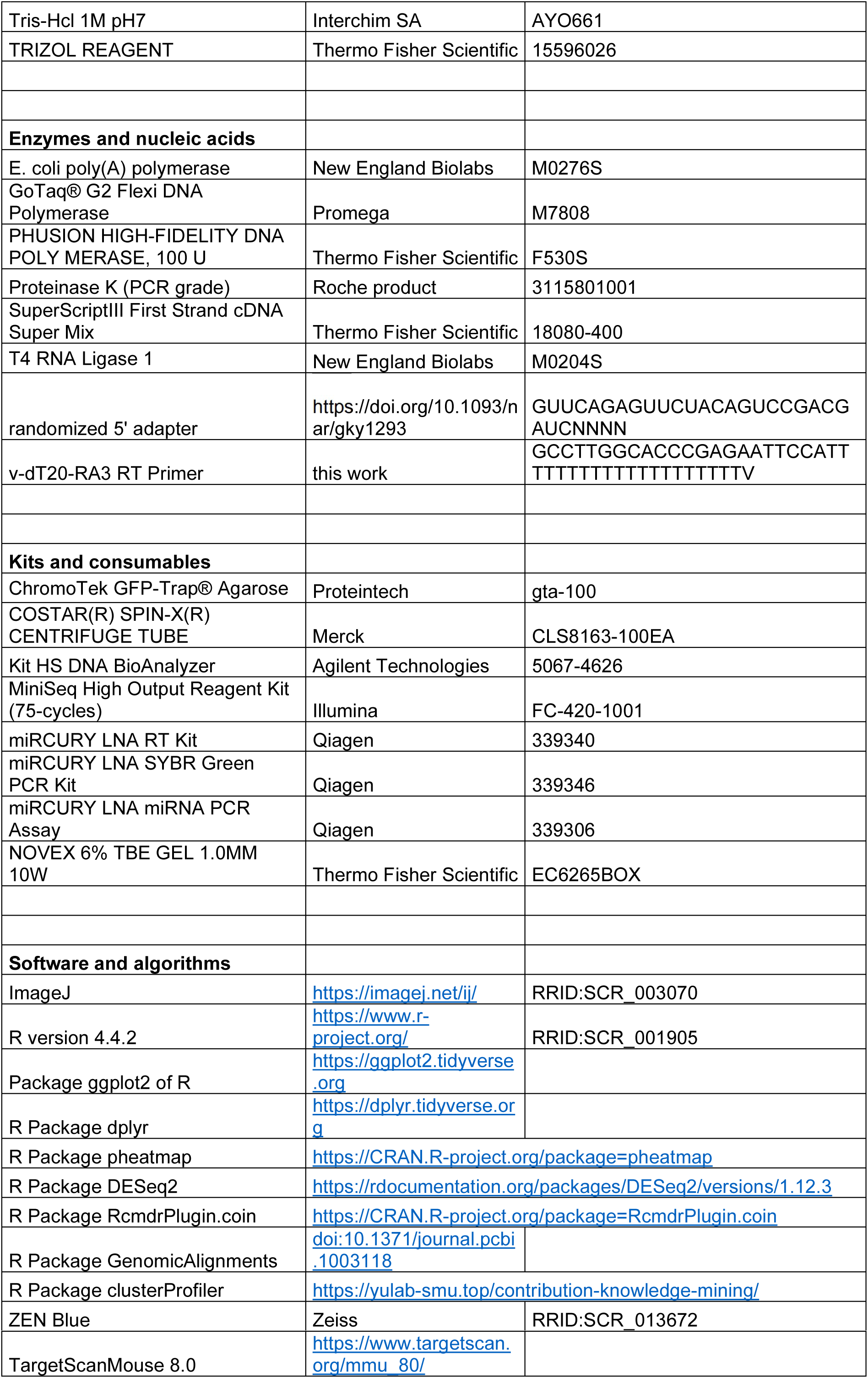

### In Vivo Electroporation and Transgenic mice

*in vivo* electroporation experiments were performed on P1 pups^70^.

The construct used to generate the T6B-FHY transgenic mouse line described in this paper was derived from the Ai14 plasmid by replacing the tomato transgene by T6B-FHY (Hauptman et al. 2015). The resulting construct was subsequently used to derive the plasmid that was introduced into the genome of the PSD95-T6B-FHY mouse line by fusing the PSD95 coding sequence to the 5’ end of T6B-FHY.

Transgenesis was performed at the SEAT transgenesis center (CNRS, Villejuif, France). The 2 constructs were integrated into the genome of 129/SvPas ES cells by homologous recombination at the ROSA-26 locus. Recombined ES cells were identified by PCR screening and subsequently microinjected into blastocytes of pseudogestant C57BL/6N females. The chimeras were then back-crossed to C57BL/6N animals.

### Brain sections preparation, immunohistochemistry for optical Imaging

Animals were deeply anesthetized with an overdose of xylasin/ketamine then intracardially perfused with 4% paraformaldehyde (w/v; PFA) in PBS. The brains were dissected out and further fixed overnight at 4°C in the same fixative. Fifty micrometer wide floating sections were processed as previously described^71^, blocked in PBS supplemented with 0.3% Triton and 5% fetal bovine serum for 1 h at room temperature and incubated at 4°C overnight in the blocking solution supplemented with primary antibody. After washing, sections were incubated for 2 h at room temperature in the blocking solution supplemented with a fluorophore-conjugated secondary antibody. Sections were mounted onto Superfrost Plus slides with Mowiol. Images were acquired using an apotome fluorescence microscope (Axioplan2, ApoTome system, Zeiss, Germany) or a laser confocal scanning microscope (LSM780, Zeiss, Germany).

### Cell Counts and Statistical Analysis

The density in fluorescent neurons was counted differently depending on the experiment. For figure 1A, the number of fluorescent neurons per OB section was counted manually. However, because this time-course experiment included multiple independent electroporation experiments performed on different litters which can cause variations in the efficiency of electroporation, we normalized the average number of fluorescent neurons per OB section with the average number of remaining fluorescent cells in the SVZ region^28^ which estimates the efficiency in electroporation. For figure 1C, the number of fluorescent neurons per OB section was counted manually. For figure 1E the number of fluorescent neurons per section was too high to be counted manually, therefore the quantity in fluorescent neurons was estimated automatically using imageJ by counting the number of fluorescent pixel. In all cases, the density was subsequently deduced by dividing the estimated number of fluorescent neurons in a given section by the section area, expressed in pixel x pixel, counted automatically with imageJ.

### Electron microscopy

Mice underwent perfusion with 4% paraformaldehyde (PFA) in phosphate-buffered saline (PBS). Following perfusion, brains were dissected out and 1 mm^2^ pieces of brain tissue were chopped. The pieces of brain were fixed with OsO4 2% in PBS for 1 hours at 4°C, then washed in PBS, dehydrated with an ethanol gradient and embedded in Epon resin. Polymerization was then performed for 48 hours at 60°C. The blocs were cut using an ultramicrotome (Leica UC7, Leica, Netherlands) to obtain 90 nm thick sections collected on Nickel grids. The grids were first treated for 1-3 minutes with a saturated solution of sodium metaperiodate, then rinsed for less than 5 minutes with 1% Triton X-100 in TBS. Next, the grids were incubated for 1 hour in 5% BSA with 0.1 % fish gelatin skin, followed by an overnight incubation with a chicken anti-GFP primary antibody (diluted 1/250 in TBS for PSD95, 1/125 for T6B). Then the grids were rinsed four times in TBS for 5 minutes, incubated 1h at 37°C in a secondary antibody (Anti-chicken coupled with 10 nm gold bead) diluted 1/30 in TBS, and washed four times in TBS for 5 minutes. Finally, the grids were treated with 2.5% Glutaraldehyde in 0.05 M CaCo for 10 minutes. Images were captured using a Transmission Electron Microscope (Tecnai G2, ThermoFisher, USA) operating at 200 kV. Imaging was facilitated by a Veleta camera (Olympus, Japan).

### AGO-APP

Olfactory bulb or cortex tissues were dissected from T6B or PSD95-T6B expressing mouse brains and immediately fixed in 1 ml 4% paraformaldehyde (PFA) for 10 minutes at 25 °C. Fixation was quenched in 245 mM glycine for 5 minutes at 25 °C and samples were washed twice with ice-cold phosphate-buffered saline (PBS) for 5 minutes at 25 °C.

Fixed tissue samples were then lysed in 1 ml of lysis buffer containing 150 mM KCl, 25 mM Tris-HCl (pH 7.5), 2 mM EDTA, supplemented freshly with 1 mM NaF, 0.5% NP-40, 1 mM DTT, and 1 mM AEBSF. Cell lysis was performed by sonication using a Vibra Cell sonicator (10 cycles of 1 minute and 20 seconds each; 10 seconds on, 10 seconds off; 34% amplitude; 4 °C). The lysate was clarified by centrifugation at 15,000 × g for 15 minutes at 4 °C.

Total protein concentration was quantified using the Pierce BCA Protein Assay Kit (Thermo Fisher Scientific, Product Number: 23227), and the lysate was adjusted to a final concentration of 2000 µg in a total volume of 1000 µl.

For each immunoprecipitation (IP), 25 µl of GFP-TRAP® Agarose (GTA) beads were used. The beads were blocked with 1% BSA for 2 hours at 4°C, followed by two washes with ice-cold PBS, and centrifuged for 2 minutes at 2500 × g. A total of 950 µl (1900 µg) of cleared lysate was added to the washed beads, and the mixture was incubated for 1 hour at 4°C with shaking or on a rotator wheel.

The beads were then washed five times with wash buffer containing 1 M NaCl, 50 mM Tris-HCl (pH 7.5), 5 mM MgCl2, supplemented with freshly added 1 mM NaF (Sigma-Aldrich, Product Number: PHR1408-1G), 0.01% NP-40, 0.1–1 mM DTT (Sigma-Aldrich, Product Number: 3483 12-3), and 0.1– 1 mM AEBSF (Sigma-Aldrich, Product Number: SBR00015-1ML). During the final wash step, the beads were transferred to a new tube precoated with PBS for at least 1 hour to prevent bead sticking. After the final wash, the beads were washed once more with ice-cold 1X PBS.

### RNA Analysis

IP samples were supplemented with 50 µl of 4% SDS in 0.1 M NaHCO₃, and incubated at 50 °C for 10 minutes with shaking at 700 rpm. The supernatant was collected, and the bead release was repeated to ensure complete recovery. Proteinase K (PCR grade, Roche, Product Number: 03115801001) was prepared at 20 mg/ml in PK buffer and pre-incubated at 37 °C for 20 minutes to inactivate potential RNases. A total of 100 µl of the prepared Proteinase K solution was added to each sample (input and IP), and the mixture was incubated overnight at 65 °C with shaking at 1000 rpm. RNA was then extracted with 1 ml of TRIzol reagent following the manufacturer protocol. At the end of the process the RNA pellets were resuspended in 12 µl of nuclease-free water.

To estimate the miRNA concentration in the AGO-APP samples a qRT-PCR was systematically performed according to the manufacturer’s instructions (Qiagen). Briefly, 3 µl of the RNA samples were reverse transcribed. The resulting complementary DNA (cDNA) was diluted 1:10, and 3 µl of the diluted cDNA was used per well for subsequent qPCR analysis using a let-7a-5p targeting probe.

### Library preparation and miRNAs Sequencing

The remaining 9 µl of the AGO-APP samples were first polyadenylated according to the manufacturer (NEB) instructions. Following enzyme deactivation (65°C for 20 minutes), RNA precipitation with ethanol and resuspension in 2.5 µl of water, a 5′ randomized adapter (Kim et al. 2019) was ligated to the polyadenylated miRNAs using T4 RNA-Ligase 1 according to manufacturer (NEB) instructions in a final volume of 10 µl. For this ligation step the adapter concentration was adapted to the amount of miRNA in the sample as estimated by the qRT-PCR previously realized. Adapting the adapter concentration allows to favor the amplification of the miRNA containing molecules. Bellow, the rule we followed:

CT of let-7a-5p 5’ adapter concentration 14-17 0.18 µM

18-19 0.05 µM

19-20 0.01 µM

21-25 0.003 µM

Subsequently 3µl of the ligation reaction was directly reverse-transcribed using the v-dT20-RA3 oligonucleotide to prime the reaction. Then the library was amplified by PCR performed on 5 µl of the ligation reaction using the Illumina primers RP1 and an RPIx index primer and following the program: 98° for 1 min - X times (98°C for 10 seconds, 58°C for 30 seconds, 72°C for 15 seconds) - 72°C for 10 min. The number X of cycles was previously determined through pilot PCR reactions performed on 1 µl of the reverse transcription reaction under the same amplification conditions but applying a variable number of cycles: 18, 21 and 25. The products of the PCR reaction were loaded on a 6% polyacrylamide TBE gel, from which the desired band of approximately 165 bp was extracted and purified. After purification, the concentration of all samples of the library was analyzed on an Agilent 2100 Bioanalyzer, according to the manufacturer’s instruction. Equal amounts of each sample were mixed for sequencing on an Illumina Miniseq instrument, according to the manufacturer’s instruction

### Bioinformatic analysis and statistics

The fastq files generated by the sequencing process were analyzed, first using the Galaxy website to trim the reads and to align the reads to the mm10 version of the mouse genome for generating the bam files, and then using the GenomicAlignments package of R to count the miRNA reads. Differential expression was analyzed using the DESeq2 package of R. To assess the enrichment in GO terms among a list of predicted target genes we used the enrichGO function of the clusterProfiler package of R. Dendrogram and heat map were generated in R using the stats and pheatmap packages, respectively. Graphs were generated in R using the ggplot2. Statistical analysis between two experimental conditions was performed with the non-parametric exact Wilcoxon test using the RcmdrPlugin.coin package in R.

